# Electrochemical Biosensor Using Polymer Nanocomposites for G6PD Deficiency Detection

**DOI:** 10.1101/2024.03.13.584424

**Authors:** Ronald Rastre Salas, Marcos Marques da Silva Paula, Yonny Romaguera Barcelay, Walter Ricardo Brito

**Affiliations:** LABEL-Laboratory of Bioelectronic and Electroanalytic, Central Analytical Lab, Federal University of Amazonas, Manaus, Amazonas, Brazil; Postgraduate Program in Materials Science and Engineering, Federal University of Amazonas, Manaus, Brazil; CEMMPRE, BioMark@UC, Department of Chemical Engineering, Faculty of Sciences and Technology, University of Coimbra, Coimbra, Portugal

**Keywords:** Biosensors, Electrochemical, G6P, DHP, D6PD, NAD^+^

## Abstract

Electrochemical biosensors based on the combination of DHP, OG, AuNPs, and G6PD, used to determine the enzymatic activity of G6PD, showed optimal current intensity values close to pH 7.8 at room temperature. Square wave voltammetry was optimized and used to evaluate the responses of the biosensors during the enzymatic reaction and construct the analytical curves. The best results were found under the following conditions: 30 minutes of incubation, in the range of NAD^+^ concentrations from 0.75 to 6 μmol L^−1^; obtaining coefficient of determination (R^2^) values between 98 and 99% about the analytical curves. The biosensors maintained their stability after five days of storage. The calculated LOD and LOQ limits showed good sensitivity of the method, achieving results in the order of 10^-6^ mol L^−1^.

## 1 INTRODUCTION

A lot of interest has been generated by biological structures for a variety of applications, including the creation of biosensors. The usage of biomolecules is appealing due to their biocompatibility and stability qualities, specificity, and capacity to detect other molecules. Examples include enzymes, antibodies, antigens, peptides, and DNA[1], [2], [3], [4]. In our situation, enzymes have attracted attention for biotransformation research and their application in industry and health as clinical diagnosticians, with a view to the creation of biosensors, due to several qualities including the possibility of separation, purification, catalytic power, and specificity [5].

Biosensors are analytical tools that can transform physical-chemical data from chemical reactions, heat release, electron transfer, pH changes, variations in protons concentration, gas release, light emission or absorption, mass variation, and oxidation state changes into a quantitative or semi-quantitative analytical signal. As a result, the materials used in the transducers must ensure that the biosensor presents a fast response in addition to being economical. The transducers, on the other hand, are responsible for converting the signal generated by the biological element into a measurable response, such as electrical current, potential, variation in temperature, and others [5].

The use of biosensors shapes their development, as do other elements that are crucial to the architecture of the sensor, such as sensitivity, sample environment, cost, life expectancy, and intended usage. As a result, the biosensor must include qualities like selectivity, which is demonstrated by the specificity of the enzymes, sampling frequency, operational stability, and repeatability of data, among others, to be employed as an analysis instrument [5], [6], [7]. The biosensor essentially consists of three integrated components: a transducer, a signal processing system, and a bioreceptor element. The sensor layer, which is made of biomolecules immobilized on the surface of a transducer and allows a concentration of the desired component contained in the sample to be measured, is the primary bioreceptor component of the biosensor. These variations, which are proportional to the analyte’s concentration, are picked up by the transducer and converted into an analytically quantifiable signal by microelectronics before being processed, amplified, and recorded [8], [9], [10]. Each sample and the sort of measurement that is of interest determine the appropriate biological material and transducer. The biocomponent controls the biosensor’s level of selectivity or specificity and detects the target substance through a chemical reaction that produces a signal. The transducer and the bioreceptor component are the two main parts of the biosensor[11].

The presence of biomolecules in biosensors is what makes them highly specific and selective devices, therefore the process of immobilizing this biomolecule on substrates enriched with carbon nanotubes, graphene, and gold nanoparticles, just to mention a few, to improve electrical conductivity becomes one of the fundamental steps when creating the biosensor. The combination with other nanomaterials has enabled the construction of biosensors for different purposes, for example: in the health sector, in the food industry where they are used for quality control and detection of food components, in the environmental area they are used to detect organophosphates and metals heavy and all thanks to the main advantages of this type of device, compared to conventional analytical methods, of being small, easy to handle, relatively cheap and performing analysis quickly and specifically [8], [12], [13].

Taking into account that, nowadays, biosensors are considered powerful analytical tools, as they make use of bio-recognition, which results in fast and sensitive responses, combining the selectivity of biochemical reactions with operational simplicity, we decided to develop a mono-enzymatic biosensor, exploiting the catalytic activity of the G6PD enzyme. The enzyme was immobilized on a gold electrode containing a polymeric network of DHP combined with gold nanoparticles (AuNPs) and graphene oxide (OG) to form three different composites DHP-AuNPs-G6PD, DHP-OG-G6PD, and DHP-OG-AuNPs-G6PD. In the system proposed here, G6P is oxidized and the NAD^+^ reduced in the catalytic reaction generates a measurable signal, where G6PD catalyzes the specific dehydrogenation of G6P consuming NAD^+^ to generate 6-phosphogluconate and NADH, a variant similar in results to that illustrated in **equation 1**.

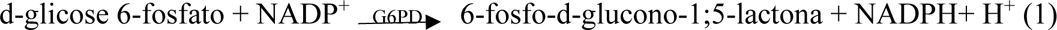

## 2. MATERIALS AND METHODS

### 2.1. Materials

Graphene oxide 2 mg mL^-1^ dispersed in water, Glucose-6-phosphate dehydrogenase 1000 U/ mL, Glucose-6-phosphate, NAD^+^ 100%, Monobasic and dibasic sodium phosphate (NaH_2_PO_4_/ Na_2_HPO_4_) obtained from Sigma-Aldrich, Sulfuric acid 98% from Neon, Isopropyl alcohol from Nuclear, KCl, Potassium ferricyanide 99% and Potassium ferrocyanide 99% from Synth, Dihexadecyl Phosphate from Sigma. All reagents used in the experiments were of analytical grade. The 70 ppm AuNPs used were prepared by the LABEL research group. Solutions were prepared with Milli-Q water with resistivity > 18 MΩ cm.

### 2.2. Equipment

To carry out the electrochemical measurements, a glass electrochemical cell with a 3-hole Teflon® lid was used to position the electrodes and add aliquots of the solutions. Disc-shaped gold electrodes, with a diameter of 2.0 mm; platinum wire and Ag/AgCl (3.0 mol L-1 KCl) were used as working, auxiliary, and reference electrodes, respectively. The electrochemical studies were carried out in potentiostats/galvonostat model PGSTAT204 (Autolab, Metrohm) coupled to a computer, using the NOVA version 2.1 program. All weighings were carried out on an analytical balance model AR2140BR (OHAUS) with an accuracy of ± 0.01 mg. pH measurements were carried out using a Five Easy F20 model pH meter (Mettler-Toledo). To stir the solutions during voltammetric measurements, a magnetic stirrer model NI1107 (Nova) was used. Homogenization of the solutions was carried out in a TS-218 ultrasound bath (Dekel). All experiments were performed at room temperature.

### 2.3. Preparation of AuE biosensors

At the end of each step, Milli-Q water was used to wash the electrode. Meticulously polish the gold electrode (AuE, 2 mm in diameter) with 0.3 and 0.05 m alumina. To remove alumina from the surface of the electrodes, the AuE was placed in an ultrasonic bath for 5 minutes while being submerged in a hydroalcoholic medium (isopropyl alcohol). At room temperature, electrochemical cleaning was carried out using 0.5 mol L^-1^ H_2_SO_4_, 0.1 Vs^-1^, and 50 cycles in the potential range of 0.0 to 1.5 V.

A mass of 1.0 mg of DHP was added to 1000 μL of 0.1 mol L^−1^ of phosphate buffer (pH 7.6) and taken to an ultrasound bath for 60 minutes in ice-cold water to obtain a dispersed solution of the DHP molecules. DHP. To 82.3 μL of this dispersion, 11.8 μL was added as a total volume of AuNPs (70 ppm), OG and the mixture (AuNPs + OG) and 5.9 μL of G6PD (5.9 U/ μL) to make room to the formation of the polymer composites under study (DHP-AuNPs-G6PD; DHP-OG-G6PD and DHP-OG-AuNPs-G6PD). These mixtures were subjected to a 30-minute ultrasound bath. Of the prepared dispersion, 2 μL was deposited on the AuE surface and allowed to dry at 25 ± 1◦C for 12 h until the solvent evaporated. **Figure 1** shows a summary of the biosensor manufacturing process.

**Figure 1.**
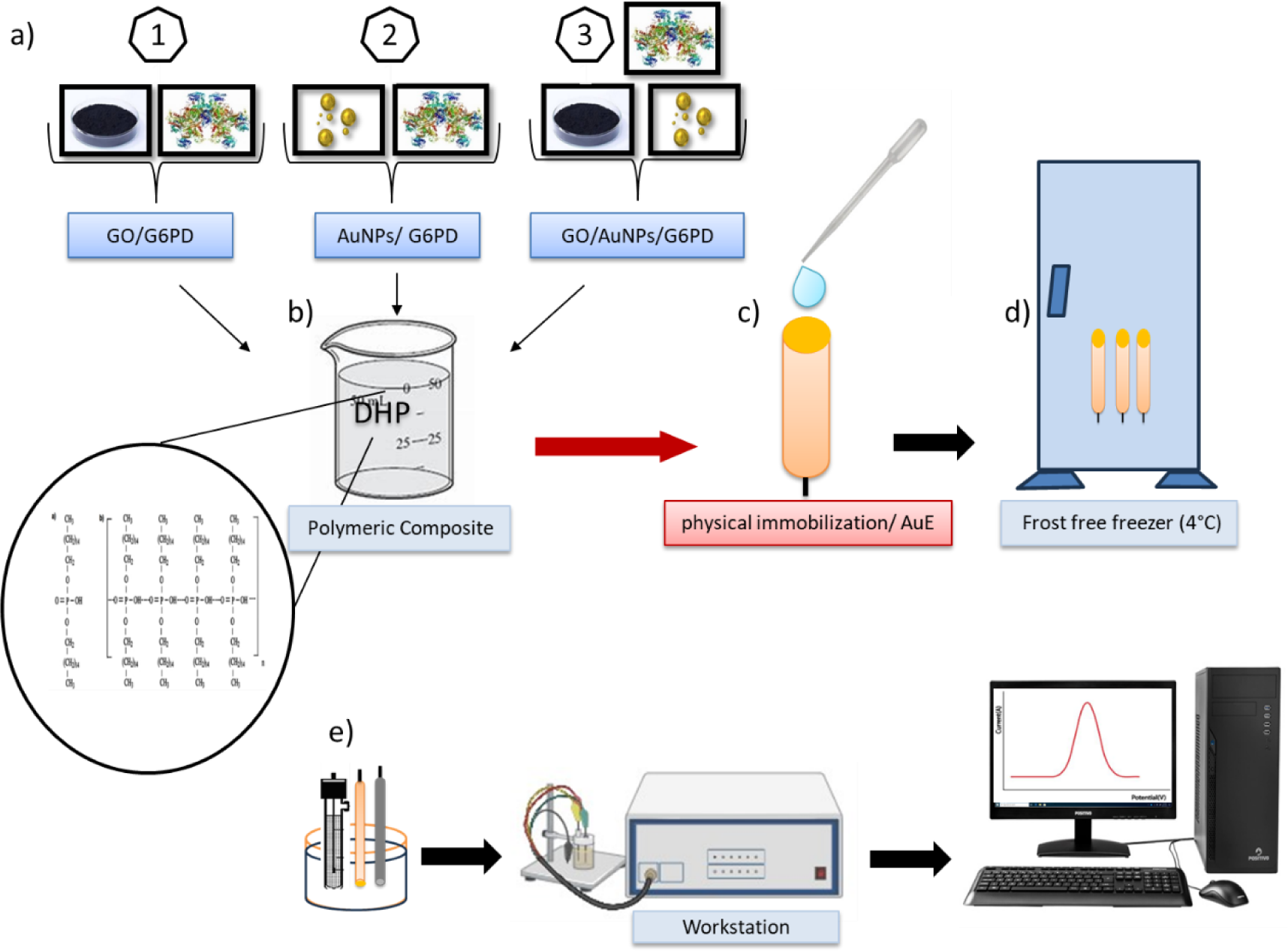
Summary of biosensor preparation.

The biosensor was not utilized and was kept at 8°C in a frost-free refrigerator.

### 2.4. Electrochemical characterization

Electrochemical evaluations were carried out for each of the steps that precede the G6PD enzyme immobilization process until reaching the biosensor itself. The electrochemical characterization study was carried out using cyclic voltammetry (VC) and electrochemical impedance spectroscopy (EIE) techniques. The VC experiments were carried out in a 0.1 mol L^−1^ PBS buffer solution containing an equimolar mixture of the salts K_3_ [Fe(CN)_6_] and K_4_ [Fe(CN)_6_] 5×10 ^−3^ mol L^−1^ in the range –0.4 to 0.8V to 0.1 V s^−1^. For EIE, a frequency in the range between 100 mHz and 100 kHz was applied at 0.1 V with a disturbance amplitude of 10 mV in a phosphate buffer solution with the same characteristics.

### 2.5. Analytical procedure

The analytical methodology was developed by carrying out a study to determine the optimal conditions for using the SWV techniques, and the results showed that for the SWV technique (frequency 25 Hz, amplitude 20 mV, and ΔE= 5 mV) in phosphate buffer 0.1 mol L-1 and pH= 7.6 using NAD^+^ (3×10-4 mol L^-1^), G6P (0.2 mol L^-1^) and Mg^2+^ (0.5 mol L^-1^) as standard solution. Responses were evaluated by the intensity of peak currents (Ip), potential displacement, and signal stability after the enzymatic reaction. As a precedent for this study, the enzymatic activity of G6PD was evaluated using a UV spectroscopy technique, determining the absorbance of the formation of the NADH molecule as a result of the reduction of the cofactor NAD^+^ in different reaction times at 340 nm. One unit of enzyme activity per milliliter (U mL^-1^) is defined as the amount of enzyme that causes an increase of 0.001 absorbance units per minute. After adding the sample, the rise in absorbance was observed for 60 minutes.

The analytical curves were built by adding repeated aliquots of various amounts of the NAD^+^ standard solution (3×10^-4^ mol L^-1^) in the electrochemical cell after the experimental parameters had been tuned. Analytical curves created in triplicate based on the rise in peak current caused by the enzyme’s catalytic activity, which is based on the oxidation of G6P and reduction of NAD^+^. Calculated on blank samples under identical experimental settings, the detection and quantification limits assess the equipment’s noise output in the potential detection ranges.

Other investigations were made, including ones into the effect of pH and the stability of the biosensor. In the first study, the 0.1 mol L^-1^ phosphate buffer solution’s pH (5.0, 6.8, 7.0, 7.6, and 8.2) was changed to determine the ideal pH for determining the enzymatic activity. In the second study, the biosensors were kept for 23 days straight while daily analyses were conducted under the same optimal conditions.

## 3. DISCUSSION OF RESULTS

### 3.1. Electrode surface characterization

According to the Randles-Sevcik **equation 2**, the electroactive area of AuE and the modification on its surface were calculated to be 0.5 mol L^-1^ KCl in the presence of 5.0 × 10 ^-6^ mol L^-1^ of [Fe(CN)_6_]^3-^ (data not shown) [14], [15].

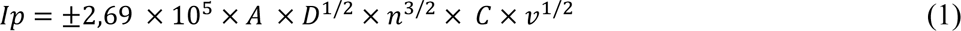

where p is the potential sweep rate (V s^-1^), n is the number of electrons transferred during the redox reaction, A is the electroactive area (cm^2^), D is the diffusion coefficient of [Fe(CN)_6_]^3-^ in solution (7.7 × 10^-6^ cm^2^s^-1^) [16], and C is the concentration of [Fe(CN)_6_]^3-^ in the bulk solution (mole cm^-3^). The electroactive areas of the biosensors (DHP-OG-G6PD; DHP-AuNPs-G6PD and DHP-OG-AuNPs-G6PD) and electrode (AuE) without modification were calculated to be 0.02770 ± 0.0002 cm2; 0.0263 ± 0.0003 cm2; 0.0270 ± 0.0006 cm^2^ and 0.03001 ± 0.0001 cm^2^ (n = 5), respectively. As demonstrated, the inclusion of the polymeric composite containing the enzyme causes a reduction in the electroactive area of the modified electrode.

### 3.2. Electrochemical characterization of AuE and modified AuE

It was necessary to carry out a preliminary study of the modified gold electrodes, considering the influence of each component on the AuE surface until reaching the final developed composite. **Figure 4** shows the cyclic voltammetry characterization of AuE and the modifications on its surface.

**Figure 4.**
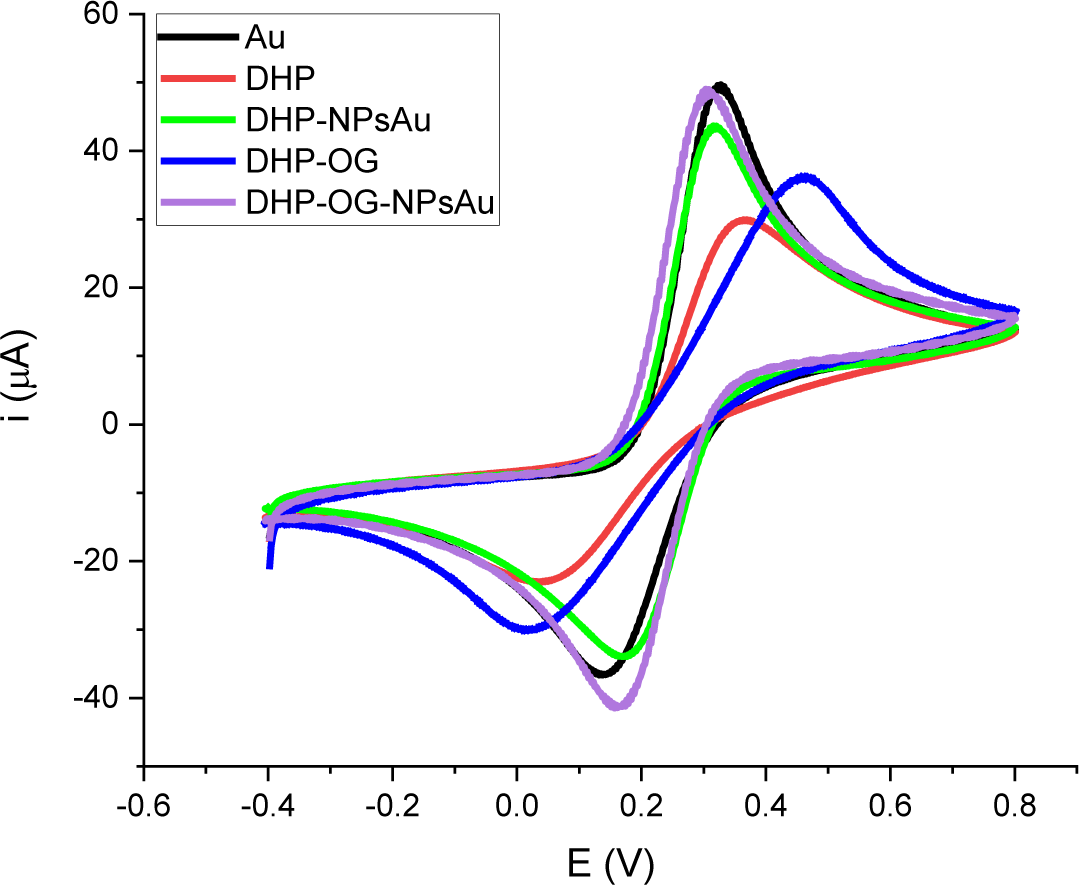
The gold electrode and the changes to its surface were electrochemically characterized by VC.

As observed in the previous figure, AuE shows a well-defined and reversible response to the faradaic currents of the redox process of the [Fe(CN)_6_] ^4–/3–^ pair, indicating that electron transfer is controlled by diffusion in the clean electrode. However, when placing the DHP film, the current intensity value (Ipa) decreases by (39.7%) and the Epa (anode peak potential) shifts to ΔEp (0.042) V due to the surfactant nature of the resulting film which slows down the charge transfer. With the addition of AuNPs, OG as well as the mixture between the two, new films (DHP-OG; DHP-AuNPs and DHP-OG-AuNPs) formed and deposited on the surface of the AuE electrode change the behavior previously described with the DHP. The films on the electrode surface show higher peak current values 36.2 < 43.6 < 49.0 µA respectively, showing how with the addition of conductive components the conductivity of the DHP film improves substantially and in the last of those presented almost like that achieved by the AuE electrode, improving the charge transfer conditions on the surface in all variants.

The previously described results are closely related to the results shown in **Figure 5**, where the EIE study stands out to investigate the electron transfer characteristics on the AuE surface and its modifications. For this study, the iron-ferricyanide redox probe (Fe(CN)_6_)^4−/3−^) was used, supported by the Nyquist graph for a better interpretation of the results of the electrode surface characteristics. As seen in the figure, the semicircular part corresponds to limited electron transfer processes and a linear part corresponds to the process favored by diffusion. The diameter of the semicircle is equal to the charge transfer resistance value (Rct) and the slope of this linear part indicates the speed at which this diffusion process is happening [17], [18].

**Figure 5.**
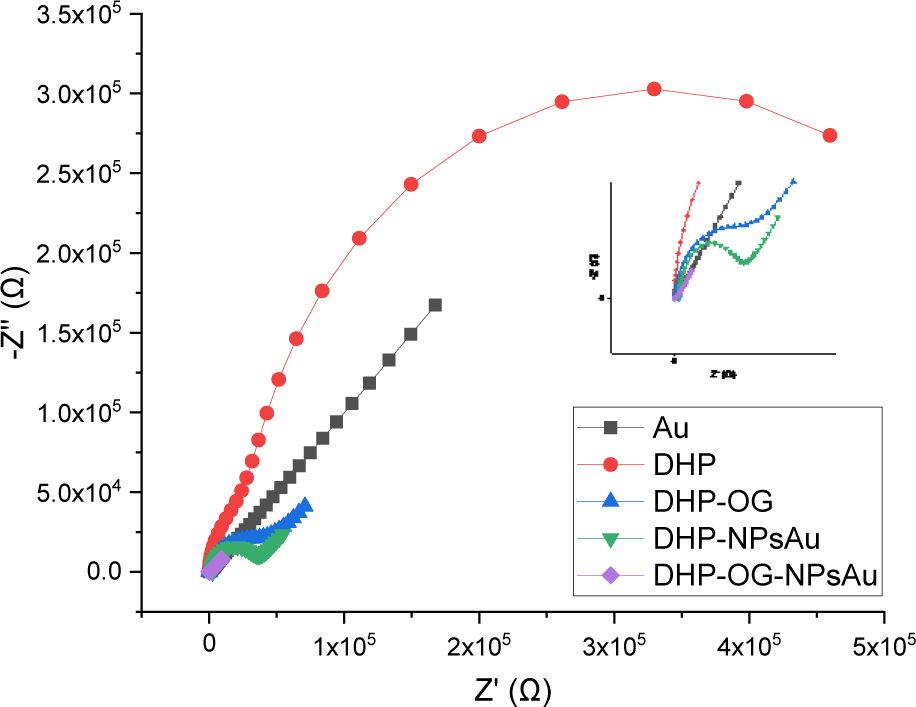
Electrochemical characterization by EIS of the gold electrode and changes on its electrode surface.

As shown in the figure, the AuE electrode has a linear profile favored by diffusion processes up to the electrode surface with charge transfer resistance (Rct) values at 501 Ω. The deposition of the DHP molecule on the electrode surface generates an Rct value of 541×103 Ω, thus generating an electrostatic blockage for the anions of the probe in use, due to the negative charge that supports the Phosphate group in the DHP molecule, which repels the interaction of probe to the electrode surface and decreases in the kinetic rate of electron transfer [19]. The films subsequently deposited on the surface provide the new modified electrode with a decrease in the value of Rct 37.8< 29.7< 7.35×103 Ω corresponding to a 98.6% maximum reduction related to the DHP-film. OG-AuNPs compared with the DHP film deposited on the surface of the AuE electrode, evidenced by the decrease in semicircles in the high-frequency region.

### 3.3. Electrochemical characterization of biosensors

In **Figure 6**, the voltammograms corresponding to the biosensors (AuE-DHP-OG-G6PD, AuE-DHP-AuNPs-G6PD AuE-DHP-OG-AuNPs-G6PD) compared with AuE as well as the EIE related to each of the systems are shown. studied. In **Figure 6A**, a decrease in anode current intensity values 11.2< 13.7< 19.7 µA is observed according to the order of appearance of the biosensors in the text, with the maximum percentage of decrease being 74.3% with the entry of the biomolecule that has bulky characteristics and blocks the surface.

**Figure 6.**
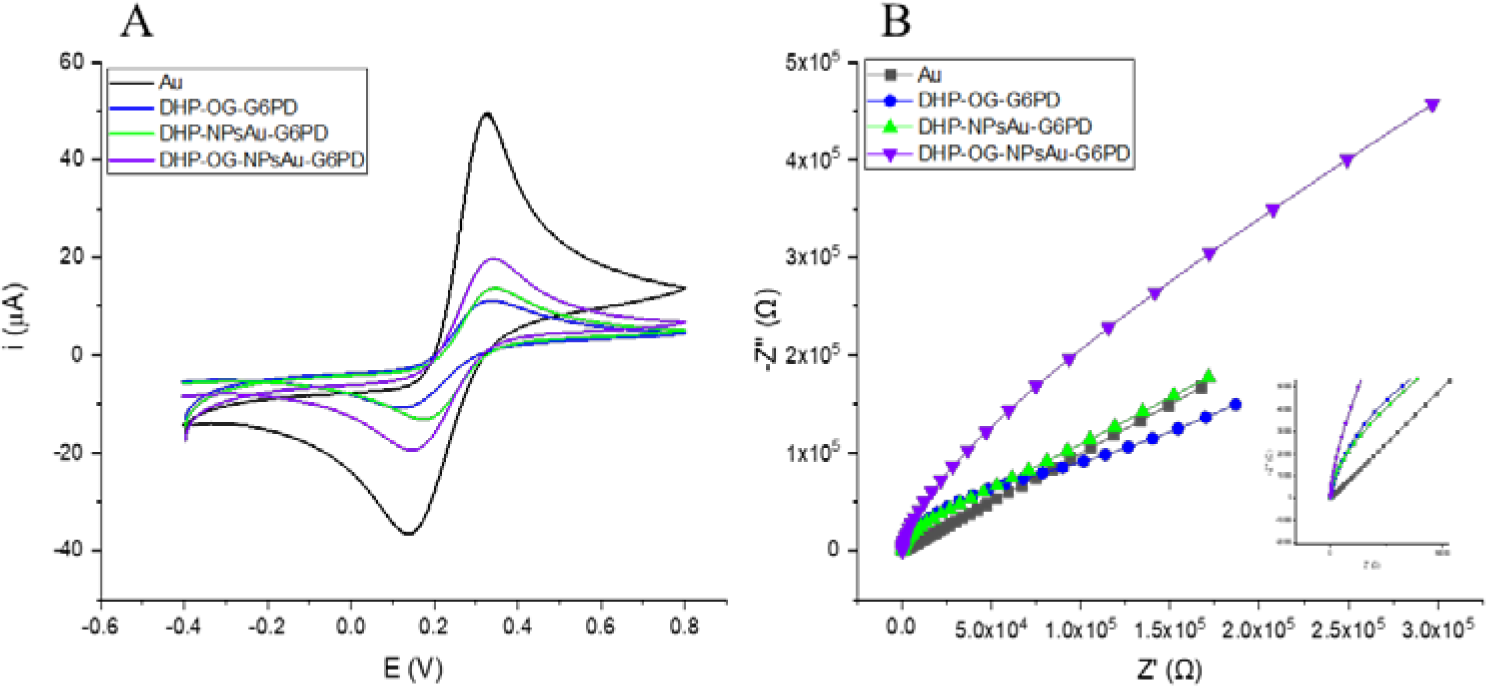
Shows VC (A) and EIS (B) electrochemical characterization.

In **Figure 6B** represented by the Nyquist graph, it confirms how, with the entry of the biomolecule, the new composite formed modifies the resistivity of the AuE surface, reaching values of Rct 105; 99.9; 50×103 Ω representing these values a considerable increase concerning the unmodified AuE.

### 3.4. G6PD catalytic activity

The determination of the enzymatic activity of G6PD and the definition of the enzymatic reaction time under the biosensor development conditions were carried out as described in the literature [20]. Triplicate measurements of the sample: 4 μL of the enzyme under study, 20 μL (G6P), 15 μL (NAD^+^), 10 μL (MgCl_2_), and made up to 2 ml with PBS solution 0.1 mol L^-1^ pH 7,6. The absorbance reading was performed at times (0, 1, 5, 10, 20, 30, 40, 50, and 60) minutes at 340 nm at room temperature. One unit of activity per milliliter (units mL^-1^) is defined as the amount of enzyme that causes an increase of 0.001 units of absorbance per minute [21], [22]. The results can be seen in **Figure 7**.

**Figure 7.** Measurement of enzymatic activity (G6PD) by reducing the cofactor NAD^+^/NADH.

The synthesis of NADH was growing over time, as shown by the rise in absorbance values, which supports G6PD’s catalytic activity. The difference between absorbance values reduces at the latter points of the curve due to a reduction in NAD^+^ molecules. The enzymatic reaction time for this study was set at 30 minutes because periods longer than this are undesirable and since various techniques for determining the activity of G6PD fall between 10 and 30 minutes. To put us on par with the fastest in the future, the research attempted to approach these times.

### 3.5. Optimization of voltametric parameters of biosensors

The square wave voltammetry technique (SWV) parameters were tuned in phosphate buffer pH= 7.6 for the biosensor before creating the analytical curves for NAD^+^. The three parameters that were modified were the frequency (10 to 91 Hz), amplitude (10 to 100 mV), and step (1 to 7 mV), with the values 25 Hz, 20, and 5 mV chosen respectively. The response was assessed using the graphic profile of the voltammogram and the current intensity data. **Figure 8** shows the corresponding voltammograms with the addition of 20 μL (G6P), 20 μL (NAD^+^), and 10 μL (MgCl_2_) in a 0.1 mol L^-1^ PBS solution and pH= 7.6 of the biosensors.

**Figure 8.**
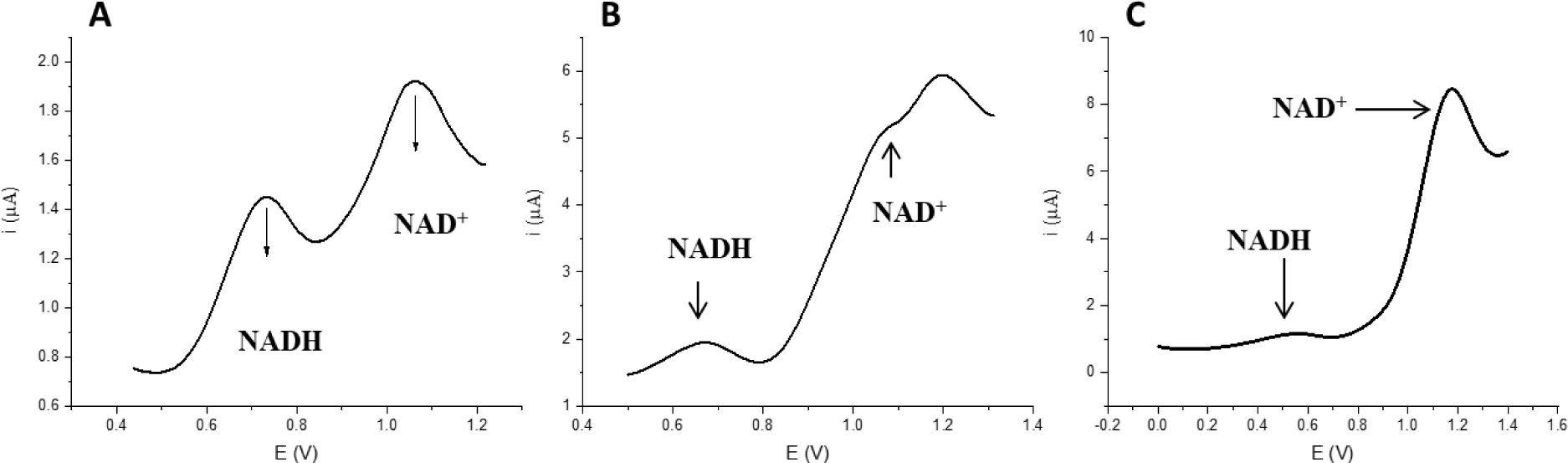
SWV of the biosensor detection process in PBS 0.1 mol L^-1^ and pH = 7.6. A (AuE-DHP-AuNPs-G6PD), B (AuE-DHP-OG-G6PD) and C (AuE-DHP-OG-AuNPs-G6PD).

**Figure 8 (A**, **B**, and **C)** shows the formation of the NADH molecule with reducing power, which is an indicator of G6PD enzymatic activity. As shown in the voltammograms in the figure under discussion where the current peaks related to NAD^+^/ NADH are identified in the potential ranges (0.82-1.3) / (0.4-0.85) V respectively. The biosensor in **Figure 8B** had the highest Ip≈ 2 µA value compared to the other biosensors. The increase in the reduction current is related to the greater formation of NADH, which is closely related to the greater number of active centers of the enzyme available to accelerate the oxidation of the G6P molecule and release the proton that is accepted by the NAD^+^ molecule.

### 3.6. Influence of Electrolyte pH on G6PD Activity

**Figure 9** displays the findings of the investigation into how the medium’s pH affects the device’s response. The studies were carried out using a standard solution of 20 L (NAD^+^) of 3× 10^-4^ mol L^-1^ in 0.1 mol PBS L^-1^ with a pH of 7.6.

**Figure 9.** Study of the influence of pH on the response of biosensors A (EBAu-DHP-AuNPs-G6PD), B (EBAu-DHP-OG-G6PD), and C (EBAu-DHP-OG-AuNPs-G6PD).

The study’s findings are displayed in the previous figure, which shows that G6PD exhibits enzymatic activity for each pH value that was investigated. However, pH=7.6 produced the best results for the current intensity. The outcome confirms previous findings in the literature that a pH level close to 7.8 is optimal for enzymatic reactions. Since the result for pH= 7.0, and 8,2 demonstrated significant peak reduction current values, it might be further examined [23], [24], [25].

### 3.7. Construction of the Analytical Curve for NAD^+^

The analytical curves in **Figure 10** were created using the standard addition method, adding aliquots of 3×10^-4^ mol L^-1^ NAD^+^ in the range of 5 to 40 µL to the electrochemical cell, stirring for 2 seconds, and incubating for 30 minutes to complete the enzyme catalysis reaction. The reading was conducted using SWV in PBS at a pH of 7.6 and 0.1 mol L^-1^.

**Figure 10.**
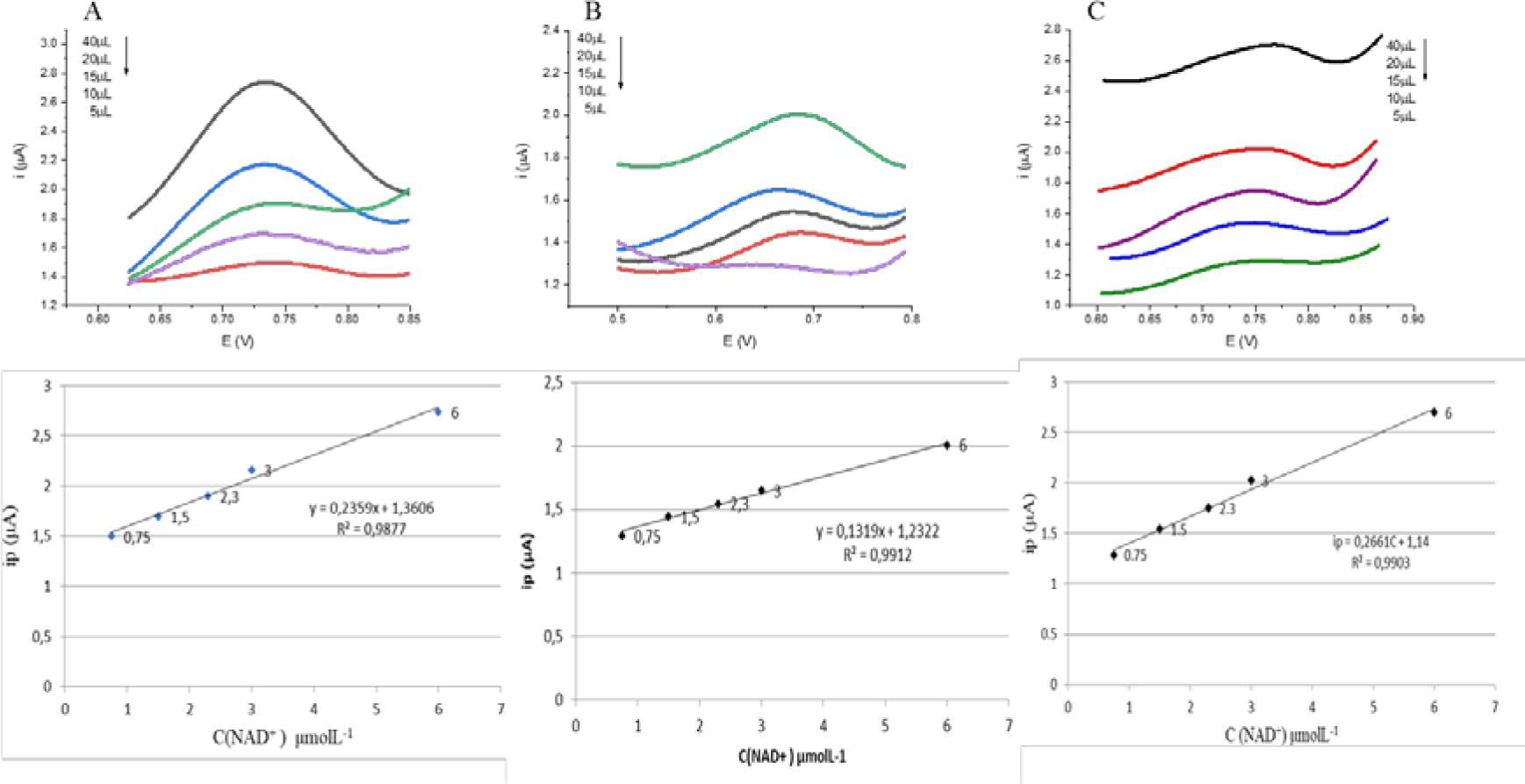
Analytical curves based on the response of the biosensors to different concentrations of NAD^+^ from 0.75 to 6 μmol L^−1^/ PBS 0.1 mol L^−1^ and pH= 7.6. A (EBAu-DHP-AuNPs-G6PD), B (EBAu-DHP-OG-G6PD), and C (EBAu-DHP-OG-AuNPs-G6PD).

**Figure 10** generally shows that there is a linear correlation between the current intensity values related to the biosensor responses and the cofactor concentration values in the solution. This linearity is evidenced in Table 1 by the straight-line equation and value of the coefficient of determination. The limits of detection (LOD) and quantification (LOQ) shown in the previously mentioned table were determined taking into account the signal/noise ratio in white samples (absence of NAD^+^) and **equations** (**2** and **3**) were used to calculate them.

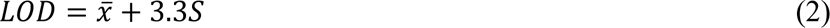

**Table 1.**
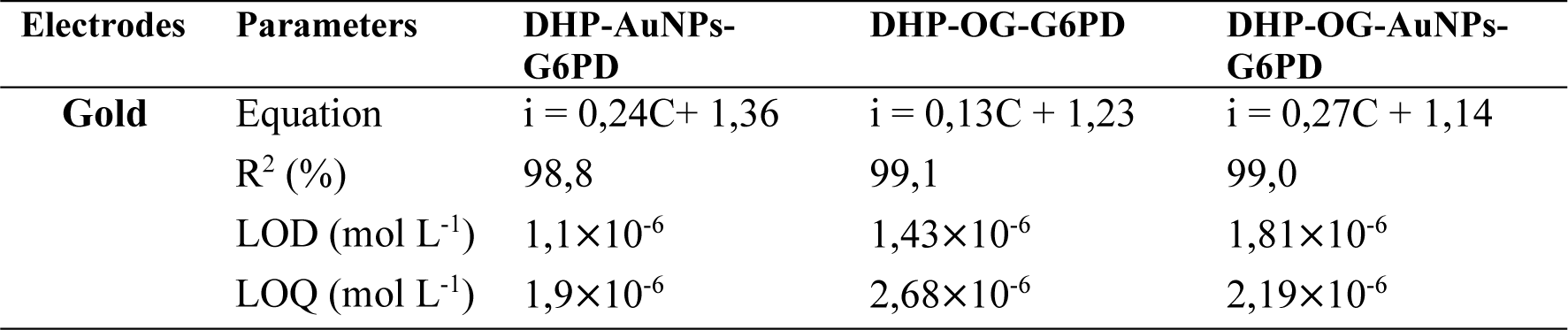
Summary of the correlation of analytical curves and LOD and LOQ.

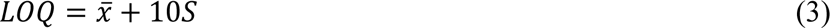

The tables shown in Tables 2 and 3 show how our results fit into a group of biosensor results reported in the literature that use G6PDH as a biological recognition element and DHP as a surface modifier.

**Table 2.**
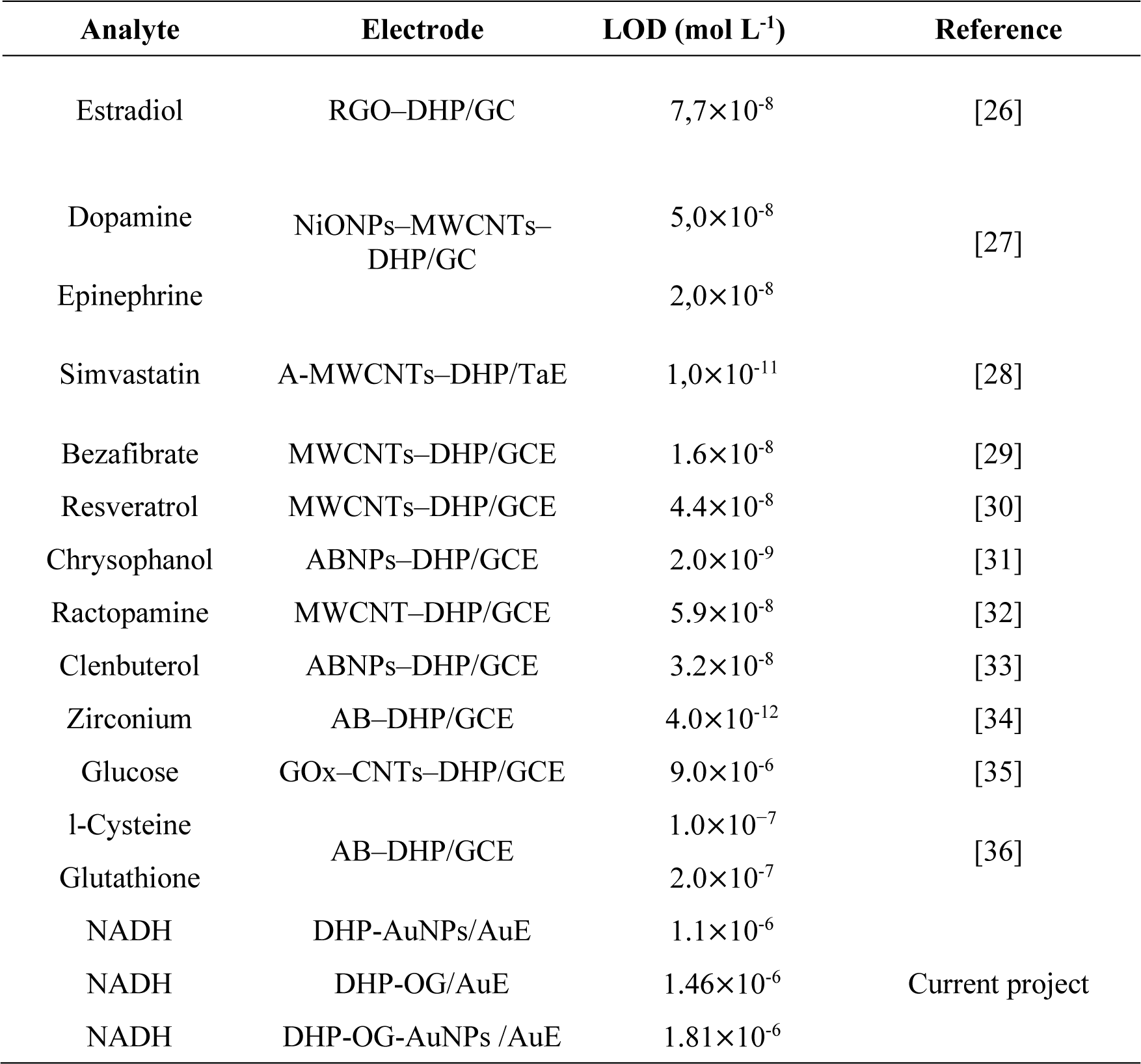
Electrodes that use DHP as a modifier.

**Table 3.**
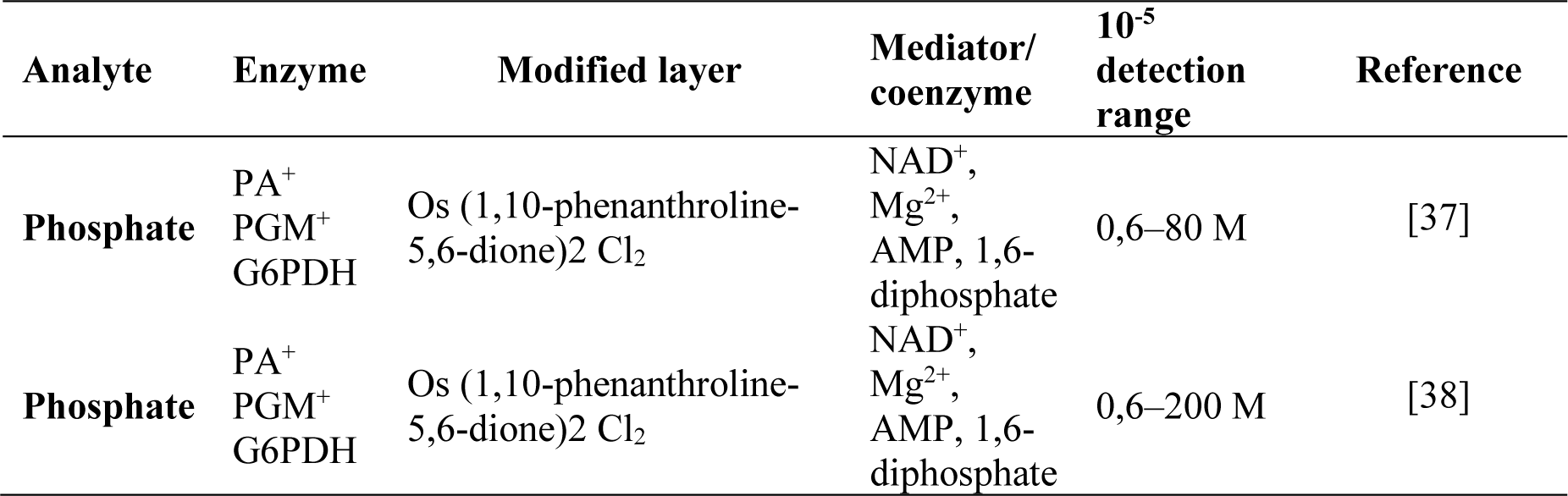
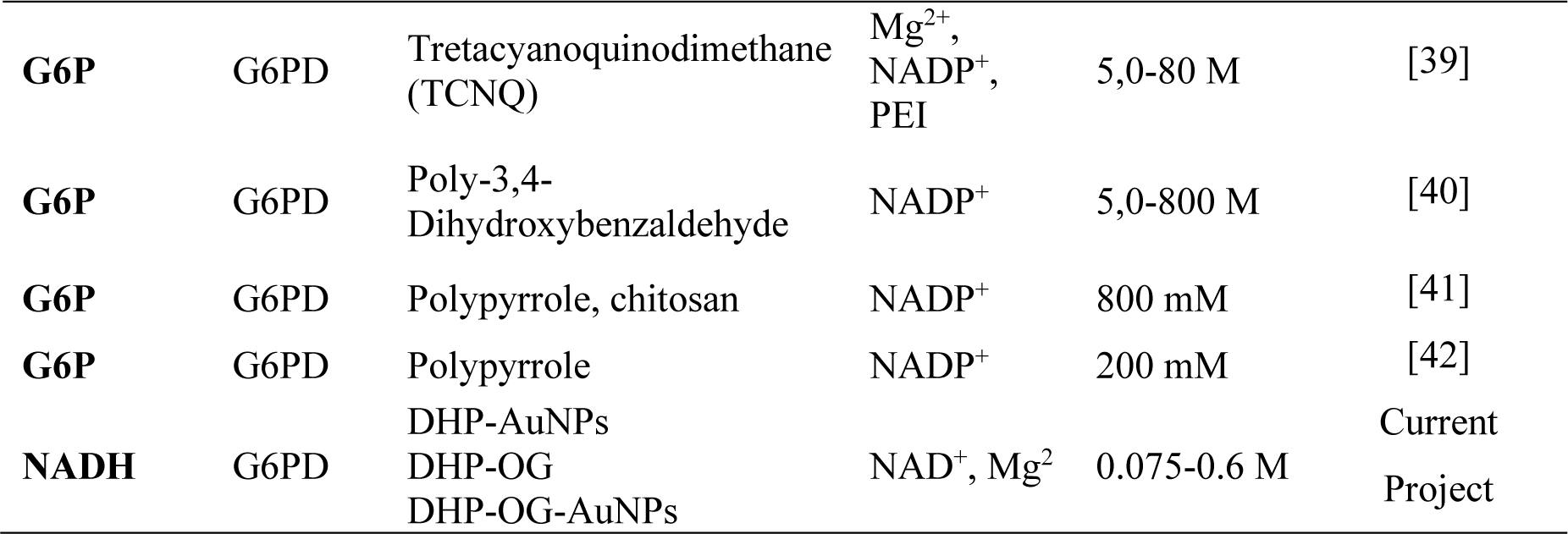
G6PD electrochemical biosensors.

### 3.8. Stability of biosensors

By examining the current values of the biosensor response following the storage time, biosensor stability experiments were carried out. **Figure 11** displays the measurements acquired during the investigation utilizing the SWV technique adjustment parameters and the addition of 20 μL of an NAD^+^ and G6P standard solution in PBS with a pH of 7.6 and 0.1 mol L^-1^.

**Figure 11.**
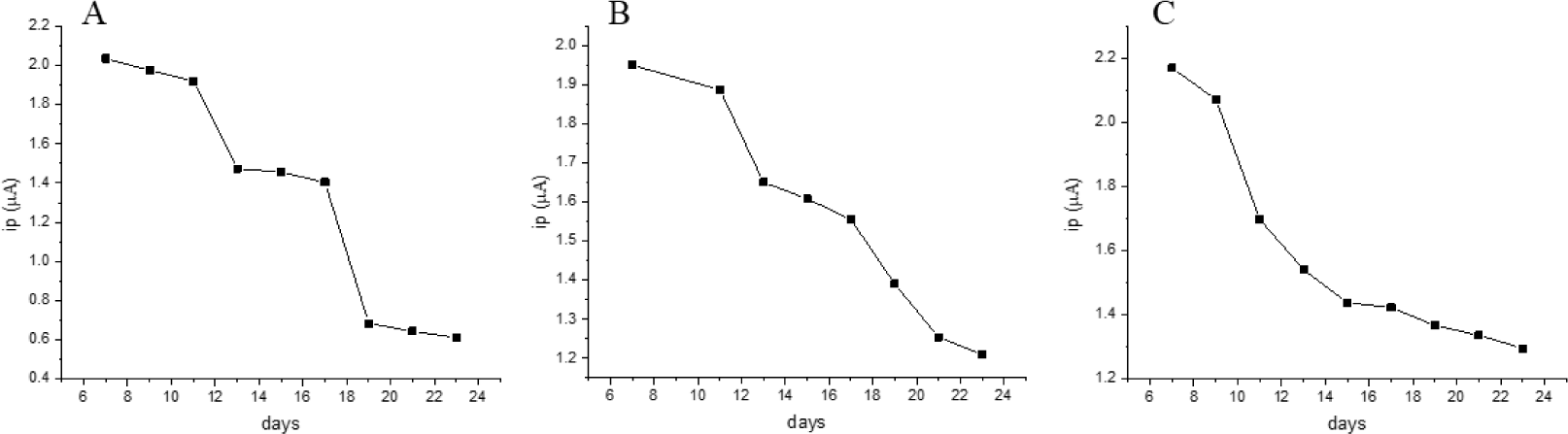
Stability study of biosensors A (EBAu-DHP-AuNPs-G6PD), B (EBAu-DHP-OG-G6PD), and C (EBAu-DHP-OG-AuNPs-G6PD).

The results in the figure shown indicated that after seven days of storage the response of the biosensors tends to gradually decrease over time, with this loss of G6PD activity being much more considerable after 23 days, recorded by the loss of current intensity values in the range 40 - 71.8%, corresponding to the following order of biosensors, EBAu-DHP-OG-G6PD, EBAu-DHP-OG-AuNPs-G6PD and EBAu-DHP-AuNPs-G6PD. Studies reported in the literature showed a significant loss of enzymatic activity within a period of 5 to 20 days [43], [44].

## 4 CONCLUSIONS

The interpretation of the results obtained electrochemically allows us to conclude that the proposed biosensors can be a tool for detecting and determining G6PD activity based on the measurement of NADH produced through enzymatic catalysis. The results of LOD (1.1; 1.43; 1.81) ×10^-6^ mol L^−1^ and LOQ (1.9; 2.19; 2.68) ×10^-6^ mol L^−1^ respectively according to the order shown in the commented figures. These results also encourage the carrying out of more advanced studies, based on the incorporation of elements that can enhance the action of the enzyme and therefore increase sensitivity as well as reduce the final cost of the biosensor. The possibility of using other G6PD immobilization techniques, e.g. the fixation of the enzyme in a sol-gel matrix, in addition to the use of different electrochemical mediators, can also be studied.

The stability study indicated that the response of the biosensors after 23 days of storage showed a tendency to decrease in the three biosensors, but EBAu-DHP-AuNPs-G6PD was most notable. The pH influence study showed that around pH 7.6 the best current intensity results were obtained, coinciding with what is said in the literature, with the EBAu-DHP-AuNPs-G6PD biosensor being highlighted in this study with the best result. Therefore, the electrochemical devices presented high analytical performance to determine the activity of G6PD, aiming at the development of a future amperometric sensor for the detection of the enzymatic deficit of G6PD in biological samples.

## ACKNOWLEDGMENTS

The authors are deeply grateful to funding institutions such as Capes, CNPQ, FAPEAM, and UFAM for providing their facilities for the development of this research and to all partner institutions.

## BIBLIOGRAPHY

[1] J. S. Graça, R. F. de Oliveira, M. L. de Moraes, and M. Ferreira, “Amperometric glucose biosensor based on layer-by-layer films of microperoxidase-11 and liposome-encapsulated glucose oxidase,” Bioelectrochemistry, vol. 96, pp. 37–42, Apr. 2014, doi: 10.1016/j.bioelechem.2014.01.001.

[2] G. Rocchitta et al., “Enzyme Biosensors for Biomedical Applications: Strategies for Safeguarding Analytical Performances in Biological Fluids,” Sensors, vol. 16, no. 6, p. 780, May 2016, doi: 10.3390/s16060780.

[3] J.-M. Moon, N. Thapliyal, K. K. Hussain, R. N. Goyal, and Y.-B. Shim, “Conducting polymer-based electrochemical biosensors for neurotransmitters: A review,” Biosens. Bioelectron., vol. 102, pp. 540–552, Apr. 2018, doi: 10.1016/j.bios.2017.11.069.

[4] E. Rosini, P. D’Antona, and L. Pollegioni, “Biosensors for D-Amino Acids: Detection Methods and Applications,” Int. J. Mol. Sci., vol. 21, no. 13, p. 4574, Jun. 2020, doi: 10.3390/ijms21134574.

[5] S. Menon, M. R. Mathew, S. Sam, K. Keerthi, and K. G. Kumar, “Recent advances and challenges in electrochemical biosensors for emerging and re-emerging infectious diseases,” J. Electroanal. Chem., vol. 878, p. 114596, Dec. 2020, doi: 10.1016/j.jelechem.2020.114596.

[6] S.-R. Yan et al., “A review: Recent advances in ultrasensitive and highly specific recognition aptasensors with various detection strategies,” Int. J. Biol. Macromol., vol. 155, pp. 184–207, Jul. 2020, doi: 10.1016/j.ijbiomac.2020.03.173.

[7] G. Mishra, A. Barfidokht, F. Tehrani, and R. Mishra, “Food Safety Analysis Using Electrochemical Biosensors,” Foods, vol. 7, no. 9, p. 141, Sep. 2018, doi: 10.3390/foods7090141.

[8] S. Singh, V. Kumar, D. S. Dhanjal, S. Datta, R. Prasad, and J. Singh, “Biological Biosensors for Monitoring and Diagnosis,” in Microbial Biotechnology: Basic Research and Applications., Nagar, India: Springer, Singapore, 2020, pp. 317–335. doi: 10.1007/978-981-15-2817-0_14.

[9] E. Cesewski and B. N. Johnson, “Electrochemical biosensors for pathogen detection,” Biosens. Bioelectron., vol. 159, p. 112214, Jul. 2020, doi: 10.1016/j.bios.2020.112214.

[10] H. H. Nguyen, S. H. Lee, U. J. Lee, C. D. Fermin, and M. Kim, “Immobilized Enzymes in Biosensor Applications,” Materials (Basel)., vol. 12, no. 1, p. 121, Jan. 2019, doi: 10.3390/ma12010121.

[11] H. Medetalibeyoglu, M. Beytur, O. Akyıldırım, N. Atar, and M. L. Yola, “Validated electrochemical immunosensor for ultra-sensitive procalcitonin detection: Carbon electrode modified with gold nanoparticles functionalized sulfur doped MXene as sensor platform and carboxylated graphitic carbon nitride as signal amplification,” Sensors Actuators B Chem., vol. 319, p. 128195, Sep. 2020, doi: 10.1016/j.snb.2020.128195.

[12] S. P. Reis et al., “Nanotubos de carbono: conceitos gerais e aplicação em biosenssores.,” Sci. Amaz., vol. 7, no. 1, pp. 53–59, 2018, [Online]. Available: http://www.scientia-amazonia.org

[13] A. Nicolia, A. Manzo, F. Veronesi, and D. Rosellini, “An overview of the last 10 years of genetically engineered crop safety research,” Crit. Rev. Biotechnol., vol. 34, no. 1, pp. 77–88, Mar. 2014, doi: 10.3109/07388551.2013.823595.

[14] M. M. Radhi, A. I. Ibrahim, Y. K. Al-Haidarie, S. A. Al-Asadi, and E. A. J. Al-Mulla, “Rifampicin: Electrochemical Effect on Blood Component by Cyclic Voltammetry Using Nano-Sensor,” Nano Biomed. Eng., vol. 11, no. 2, pp. 150–156, May 2019, doi: 10.5101/nbe.v11i2.p150-156.

[15] G. Ibáñez-Redín, T. A. Silva, F. C. Vicentini, and O. Fatibello-Filho, “Effect of carbon black functionalization on the analytical performance of a tyrosinase biosensor based on glassy carbon electrode modified with dihexadecylphosphate film,” Enzyme Microb. Technol., vol. 116, pp. 41–47, Sep. 2018, doi: 10.1016/j.enzmictec.2018.05.007.

[16] W. S. Fernandes-Junior et al., “Electrochemical Sensor Based on Nanodiamonds and Manioc Starch for Detection of Tetracycline,” *J*. Sensors, vol. 2021, pp. 1–10, Mar. 2021, doi: 10.1155/2021/6622612.

[17] Y. Chen, G. Zheng, Q. Shi, R. Zhao, and M. Chen, “Preparation of thiolated calix[8]arene/AuNPs/MWCNTs modified glassy carbon electrode and its electrocatalytic oxidation toward paracetamol,” Sensors Actuators B Chem., vol. 277, pp. 289–296, Dec. 2018, doi: 10.1016/j.snb.2018.09.012.

[18] J. Park et al., “Ultrasensitive detection of fibrinogen using erythrocyte membrane-draped electrochemical impedance biosensor,” Sensors Actuators B Chem., vol. 293, pp. 296–303, Aug. 2019, doi: 10.1016/j.snb.2019.05.016.

[19] C. Zhu et al., “An Amperometric Biomedical Sensor for the Determination of Homocysteine Using Gold Nanoparticles and Acetylene Black-Dihexadecyl Phosphate-Modified Glassy Carbon Electrode,” Micromachines, vol. 14, no. 1, p. 198, Jan. 2023, doi: 10.3390/mi14010198.

[20] D. L. Nelso and M. M. Cox, Princípios de Bioquímica de Lehninger, 8th ed. Porto Alegre: Artmed Editora, 2022.

21. R. Rastre, M. M. da S. de Paula, and W. R. Brito, “Electrochemistry biosensor for enzymatic activity of G6PD,” Eur. Acad. Res., vol. 4, no. 12, pp. 7004–7016, 2022.

[22] D. P. C. De Oliveira, F. W. P. Ribeiro, H. Becker, P. Lima-Neto, and A. N. Correia, “Biossensor eletroquímico baseado na enzima tirosinase para a determinação de fenol em efluentes,” Quim. Nova, vol. 38, no. 7, pp. 924–931, 2015, doi: 10.5935/0100-4042.20150086.

[23] A. Ishaque, M. Milhausen, and H. R. Levy, “On the absence of cysteine in glucose 6-phosphate dehydrogenase from Leuconostocmesenteroides,” Biochem. Biophys. Res. Commun., vol. 59, no. 3, pp. 894–901, Aug. 1974, doi: 10.1016/S0006-291X(74)80063-X.

[24] C. Olive and H. Richard Levy, “[44] Glucose-6-phosphate dehydrogenase from Leuconostoc mesenteroides,” in Methods Enzymol, vol. 41, 1975, pp. 196–201. doi: 10.1016/S0076-6879(75)41046-1.

[25] P. Rowland, A. K. Basak, S. Gover, H. R. Levy, and M. J. Adams, “The three– dimensional structure of glucose 6–phosphate dehydrogenase from Leuconostoc mesenteroides refined at 2.0 Å resolution,” Structure, vol. 2, no. 11, pp. 1073–1087, Nov. 1994, doi: 10.1016/S0969-2126(94)00110-3.

[26] B. C. Janegitz, F. A. dos Santos, R. C. Faria, and V. Zucolotto, “Electrochemical determi-nation of estradiol using a thin film containing reduced graphene oxide and dihexadecylphosphate,” Mater. Sci. Eng. C, vol. 37, no. 1, pp. 14–19, 2014, doi: 10.1016/j.msec.2013.12.026.

[27] L. C. S. Figueiredo-Filho, T. A. Silva, F. C. Vicentini, and O. Fatibello-Filho, “Simultaneous voltammetric determination of dopamine and epinephrine in human body fluid samples using a glassy carbon electrode modified with nickel oxide nanoparticles and carbon nanotubes within a dihexadecylphosphate film,” Analyst, vol. 139, pp. 2842–2849, 2014, doi: 10.1039/C4AN00229F.

[28] H. Fayazfar, A. Afshar, A. Dolati, and M. Ghalkhani, “Modification of well-aligned carbon nanotubes with dihexadecyl hydrogen phosphate: application to highly sensitive nanomolar detection of simvastatin,” J. Appl. Electrochem., vol. 44, no. 2, pp. 263–277, 2014, doi: 10.1007/s10800-013-0654-y.

[29] J. A. Ardila, G. G. Oliveira, R. A. Medeiros, and O. Fatibello-Filho, “Square-wave adsorptive stripping voltammetric determination of nanomolar levels of bezafibrate using a glassy carbon electrode modified with multi-walled carbon nanotubes within a dihexadecyl hydrogen phosphate film,” Analyst, vol. 139, pp. 1762–1768, 2014, doi: 10.1039/c3an02016a.

[30] W. Huang, S. Luo, D. Zhou, S. Zhang, and K. Wu, “Electrochemical determination of resveratrol using multi-walled carbon nanotubes-modified glassy carbon electrode,” Nanosci. Nanotechnol. Lett., vol. 5, no. 3, pp. 367–371, 2013, doi: 10.1166/nnl.2013.1542.

[31] Y. Zhang, Y. Wang, K. Wu, S. Zhang, Y. Zhang, and C. Wan, “Electrochemical determination of chrysophanol based on the enhancement effect of acetylene black nanoparticles,” vol. 103, pp. 94–98, 2013, doi: 10.1016/j.colsurfb.2012.10.015.

[32] Z. Liu, Y. Zhou, Y. Wang, Q. Cheng, and K. Wu, “Enhanced oxidation and detection of toxic ractopamine using carbon nanotube film-modified electrode,” Electrochim. Acta, vol. 74, pp. 139–144, 2012, doi: 10.1016/j.electacta.2012.04.041.

[33] G. Fan et al., “Enhanced oxidation and detection of toxic clenbuterol on the surface of acetylene black nanoparticle-modified electrode,” vol. 169, pp. 102– 105, 2012, doi: 10.1016/j.molliq.2012.02.013.

[34] P. Deng, Y. Feng, and J. Fei, “Trace determination of zirconium by adsorptive anodic stripping voltammetry of its complex with alizarin violet using a glassy carbon electrode modified with acetylene black–dihexadecyl hydrogen phosphate composite film,” Microchem. Acta, vol. 175, pp. 233–240, 2011, doi: 10.1007/s00604-011-0690-4.

[35] B. C. Janegitz, R. Pauliukaite, M. E. Ghica, C. M. A. Brett, and O. Fatibello-Filho, “Direct electron transfer of glucose oxidase at glassy carbon electrode modified with functionalized carbon nanotubes within a dihexadecylphosphate film,” Sensors Actuators B Chem., vol. 158, no. 1, pp. 411–417, 2011, doi: 10.1016/j.snb.2011.06.048.

[36] L. Lin et al., “LC-ED with an acetylene black-dihexadecyl hydrogen phosphate composite film-modified electrode for in vivo analysis of thiols in rat striatal microdialysate,” Chromatographia, vol. 72, no. 5–6, pp. 447–452, 2010, doi: 10.1365/s10337-010-1682-y.

[37] J. Parellada et al., “Imunossensores amperométricos e eletrodos de enzima para aplicações ambientais,” Anal. Chim. Acta, vol. 362, no. 1, pp. 47–57, 1998.

[38] J. J. Fernández, J. R. López, X. Correig, and I. Katakis, “Biossensores de fosfato de pasta de carbono sem reagente: estudos preliminares,” Sensores Actuators B, vol. 47, pp. 13–20, 1998.

[39] A. Bassi, D. Tang, and M. Bergougnou, “Biossensor amperométrico mediado para monitoramento de gucose-6-fosfato com base em glicose-6-fosfato desidrogenase, íons Mg 2+, tetracianoquinodimetano e nicotinamida adenina dinucleotídeo fosfato em pasta de carbono.,” Anal. Biochem, vol. 268, no. 2, pp. 223–228, 1999, doi: 10.1006/abio.1998.3082.

[40] C. H. Tzang, R. Yuan, and M. Yang, “Biossensores voltamétricos para a determinação de formato e glicose-6-fosfato com base na medição de NADH e NADPH gerados por desidrogenase,” Biossensores e bioeletrônica, vol. 16, no. 3, pp. 211–219, 2001, doi: 10.1016/S0956-5663(00)00143-3Obtenha direitos e conteúdo.

[41] S. Sahin, I. Ozmen, G. Y. Bastemur, and S. P. Ozkorucuklu, “Development of Voltammetric Glucose-6-phosphate Biosensors Based on the Immobilization of Glucose-6-phosphate Dehydrogenase on Polypyrrole- and Chitosan-Coated Fe3O4 Nanoparticles/Polypyrrole Nanocomposite Films,” Appl. Biochem. Biotechnol., vol. 188, no. 4, pp. 1145–1157, 2019, doi: 10.1007/s12010-019-02979-2.

[42] S. Selmihan, O. Ismail, Y. B. Gizem, and S. P. Ozkorucuklu, “Development of Voltammetric Glucose-6-phosphate Biosensors Based on the Immobilization of Glucose-6-phosphate Dehydrogenase on Polypyrrole- and Chitosan-Coated Fe 3 O 4 Nanoparticles/Polypyrrole Nanocomposite Films,” Appl. Biochem. Biotechnol., vol. 188, no. 4, pp. 1145–1157, 2019, doi: 10.1007/s12010-019-02979-2.

[43] A. S. Bassi, D. Tang, and M. A. Bergougnou, “Mediated, Amperometric Biosensor for Glucose-6-Phosphate Monitoring Based on Entrapped Glucose-6-Phosphate Dehydrogenase, Mg2+Ions, Tetracyanoquinodimethane, and Nicotinamide Adenine Dinucleotide Phosphate in Carbon Paste,” Anal. Biochem., vol. 268, no. 2, pp. 223–228, Mar. 1999, doi: 10.1006/abio.1998.3082.

[44] Y. Cui, J. P. Barford, and R. Renneberg, “Amperometric determination of phosphoglucomutase activity with a bienzyme screen-printed biosensor,” Anal. Biochem., vol. 354, no. 1, pp. 162–164, Jul. 2006, doi: 10.1016/j.ab.2006.03.045.

